# hsegHMM: Hidden Markov Model-based Allele-specific Copy Number Alteration Analysis Accounting for Hypersegmentation

**DOI:** 10.1101/410845

**Authors:** Hyoyoung Choo-Wosoba, Paul S Albert, Bin Zhu

## Abstract

**Background:** Somatic copy number alternation (SCNA) is a common feature of the cancer genome and is associated with cancer etiology and prognosis. The allele-specific SCNA analysis of a tumor sample aims to identify the allele-specific copy numbers of both alleles, adjusting for the ploidy and the tumor purity. Next generation sequencing platforms produce abundant read counts at the base-pair resolution across the exome or whole genome which is susceptible to hypersegmentation, a phenomenon where numerous regions with very short length are falsely identified as SCNA.

**Results:** We propose hsegHMM, a hidden Markov model approach that accounts for hypersegmentation for allele-specific SCNA analysis. hsegHMM provides statistical inference of copy number profiles by using an effcient E-M algorithm procedure. Through simulation and application studies, we found that hsegHMM handles hypersegmentation effectively with a t-distribution as a part of the emission probability distribution structure and a carefully defined state space. We also compared hsegHMM with FACETS which is a current method for allele-specific SCNA analysis. For the application, we use a renal cell carcinoma sample from The Cancer Genome Atlas (TCGA) study.

**Conclusions:** We demonstrate the robustness of hsegHMM to hypersegmentation. Furthermore, hsegHMM provides the quantification of uncertainty in identifying allele-specific SCNAs over the entire chromosomes. hsegHMM performs better than FACETS when read depth (coverage) is uneven across the genome.

## Background

Characterizing somatic copy number alterations (SCNAs) is important for understanding tumorgenesis ([1]), cancer etiology and prognosis ([2]). In normal cells, two copies of chromosome are inherited from both parents. In contrast, tumor cells frequently contain alterations in copy numbers across the chromosomes, such as deletions, insertions, or amplifications among others ([3, 4]). In addition, tumor tissues always contain normal cells (reduced tumor purity) and frequently show an abnormal number of chromosomes (aneuploidy). These characteristics of the cancer genome and tissue heterogeneity complicate the estimation of SCNAs, in contrast to germline copy number variations (CNVs) analysis where neither should be considered ([5, 6]). Allele-specific SCNA analysis estimates the integer copy number for each allele instead of the total copy number, and is essential to identify the copy-neutral loss of heterozygosity (NLOH) ([7, 8]). Based on the total copy numbers only, NLOH will be misidentified as normal regions with the copy number two, when one chromosome is duplicated but the corresponding homologous region is deleted ([9]).

In this paper, we consider next-generation sequencing (NGS) platform-based whole exome sequencing (WES) data for studying SCNAs. The NGS technology provides high resolution at the single base-pair, which comes with mapping bias and the tendency for hypersegmentation. Mapping bias occurs from higher mapping rates for the reference allele than those for the variant allele at heterozygous loci([10]). This bias leads to incorrect interpretations of allele-specific SCNAs. Hypersegmentation is also a major challenge in NGS-based allele-specific SCNA. The quality of such data depends on the sample preparation, the library preparation, and polymerase chain reaction (PCR) techniques from applying NGS technology, and the exome enrichment platforms from WES. It has been reported that capture effciency could vary across the percentage of guanine or cytosine contained in DNA [11]. These technical procedures have a limit of accurate quantification of sequences, which potentially increase measurement errors that in turn result in excessive segmentations.

A number of papers have been proposed to address these challenges and complexities. While PennCNV ([12]) and QuantiSNP ([13]) are based on the assumption of 100% tumor purity, ASCAT ([14]), GPHMM ([5]), and MixHMM ([15]) account for both the tumor purity and the ploidy. However, these methods do not explicitly characterize the genotype at each allelic location. Furthermore, these methods use a B-allele frequency (BAF), which is sensitive to mapping bias. Shen and Seshan ([16]) developed FACETS that uses log Odds Ratio (logOR) instead of BAF, since the logOR of tumor versus normal cells provides unbiased allelic information. FACETS uses a genotype mixture model, providing an allele-specific tumor copy number profile adjusted for the tumor purity and ploidy. However, since the segmentation and genotype mixture modeling are conducted by separate algorithms, it is not possible to assess uncertainty in the estimation of allele-specific SCNA.

In this paper, we propose a novel hidden Markov modeling approach (hsegHMM) for allele-specific SCNA analysis accounting for the hypersegmentation. hsegHMM embeds logOR and logR (log R ratio) into a hidden Markov model (HMM) framework, and simultaneously conducts the segmentation and genotype mixture modeling required to identify SCNAs across chromosomes. Similar to FACETS, the logOR is applied instead of the BAF to adjust for mapping bias. Hypersegmentation, which may result from logR outliers, is accounted for by assuming a t-distribution for the distribution of logR. hsegHMM makes inference about allele-specific SCNAs using the E-M algorithm where we iterate between the E- and the M-step until convergence. The E-step is made tractable by using a recursive forward-backward algorithm that evaluates functions of the hidden locus-specific genotype states given the observed logR and logOR. Given a genotype state, the tumor purity and the ploidy are obtained in the M-step by maximizing the expectation of the conditional log-likelihood function.

We apply hsegHMM to a renal carcinoma cell dataset (TCGA-KL-8331) from TCGA (the Cancer Genome Atlas) project (https://cancergenome.nih.gov) to show the effectiveness of the proposed HMM framework. We also provide various simulation studies to show that hsegHMM is able to accurately detect genotype status across chromosomes that exhibit hypersegmentation. Further, through analysis and simulations, we compare hsegHMM with FACETS.

## Methods

### Hidden Markov Model:hsegHMM

#### Modeling with logRatio and logOdds Ratio as outcomes

Let *Z_k_* be the unknown genotype as the hidden state of the k*^th^* genomic location and *W_k_* and *X_k_* be the corresponding logR and logOR, respectively. We specify 12 different states of *Z_k_*, given in Table 1. The index for hidden states, *j* is from 1 to *J* (in this paper, *J* = 12) and the index for genomic locations, *k* is from 1 to *N*. The expectations of logR and logOR are given in Van Loo et al. ([14]) and Shen and Seshan ([16]), respectively. Specifically, we can write

**Table 1.** Description of tumor genotype states and corresponding genotype of total copy number: homozygous deletion (HOMD), hemizygous deletion LOH (DLOH), copy neutral LOH (NLOH), diploid heterozygous (HET), gain of 1 allele (GAIN), amplified LOH (ALOH), allele-specific copy number amplification (ASCNA), balanced copy number amplification (BCNA), and unbalanced copy number amplification (UBCNA).

| State ($j$) | Genotype | Copy Number ($C_T$) | Allelic information |
| --- | --- | --- | --- |
| 1 | 0 | 0 | HOMD |
| 2 | A | 1 | DLOH |
| 3 | AA | 2 | NLOH |
| 4 | AB | 2 | HET |
| 5 | AAB | 3 | GAIN |
| 6 | AAA | 3 | ALOH |
| 7 | AAAB | 4 | ASCNA |
| 8 | AABB | 4 | BCNA |
| 9 | AAAA | 4 | ALOH |
| 10 | AAAAB | 5 | ASCNA |
| 11 | AAABB | 5 | UBCNA |
| 12 | AAAAA | 5 | ALOH |

**Table 2.** Summary of hsegHMM-N, hsegHMM-T, and hsegHMM-TA/AB models of a renal cell carcinoma sample: hsegHMM-TA/AB indicates the hsegHMM-T with A and AB state space; Est and logL represent estimated values for parameters and log-likelihood function values given all the estimates, respectively; *Ψ* is the ploidy and *α* is the tumor purity; *κ*^2^ is the variance component of logR in hsegHMM-T; *V* (*W*) and *τ*^2^ are the variance of logR and logOR in both models, respectively; SE_H_ indicates the average asymptotic standard errors of estimates based on the Hessian matrices.

|  | hsegHMM-N |  | hsegHMM-T |  | hsegHMM-T<sub>A/AB</sub> |  |
| --- | --- | --- | --- | --- | --- | --- |
|  | Est | SE<sub>H</sub> | Est | SE<sub>H</sub> | Est | SE<sub>H</sub> |
| $\psi$ | 1.62 | 0.003 | 1.61 | 0.003 | 1.60 | 0.003 |
| $\alpha$ | 0.87 | 0.002 | 0.88 | 0.002 | 0.88 | 0.002 |
| $\kappa^2$ | N/A | | 0.16 | 0.002 | 0.17 | 0.003 |
| $V(W)$ | 0.25 | 0.002 | 0.26<sup>a</sup> | 0.003<sup>b</sup> | 0.27<sup>a</sup> | 0.003<sup>b</sup> |
| $\tau^2$ | 0.57 | 0.012 | 0.58 | 0.012 | 0.57 | 0.014 |
| $v$ | N/A | | 5.50 | 0.185 | 5.48 | 0.218 |
| AIC | 64682.17 |  | 63120.48 |  | 62923.90 |  |
| BIC | 65934.07 |  | 64380.89 |  | 62992.03 |  |

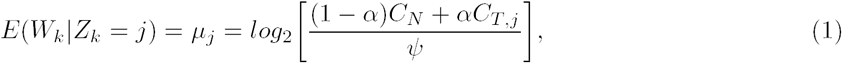

where *C_N_* is the copy number of normal cells prespecified as *C_N_* = 2; *C_T,j_* is the copy number of tumor cells at the *j^th^* state; *α* is the tumor purity proportion of the tumor tissue over a range of 0 to 1; *Ψ* is the ploidy. For example, if a tumor sample contains 100 % tumor cells and is diploidy, then *μ*_3_, the expectation of logR at the state *j* = 3 is 0 along with *C*_*T,*3_ = 2, *Ψ* = 2, and *α* = 1.

The expectation of logOR is given by

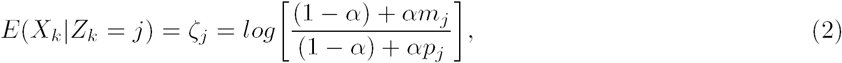

where *m_j_* and *pj* are the maternal and paternal copy numbers of the tumor at the *k^th^* genomic location, respectively. We assume that ***Z*** = {*Z*_1_, *Z*_2_, *Z*_3_, ·· ·, *Z_N_* }, the genotype sequence across chromosomes follows a Markov chain with a transition probability, *P_ij_* = *P* (*Z_k_* = *j*|*Z*_*k-*1_ = *i*) and an initial probability, *r*_0*j*_ = *P* (*Z*_1_ = *j*). *P_ij_* indicates the probability that the *j^th^* genotype state occurs conditionally on the *i^th^* genotype state at the previous location.

#### Joint emission probability with conditional distributions of logR and logOR

We consider a t-distribution for logR in order to account for outliers that are apparent in NGS data and potentially lead to hypersegmentation. Following Peel and McLachlan ([17]), we specify a t-distribution for *W_k_* with a degree of freedom *v* by a mixture of a normal distribution *N*(*W_k_*|*μ_j_*, *κ*^2^/*u_k_*) with a gamma distribution 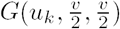,

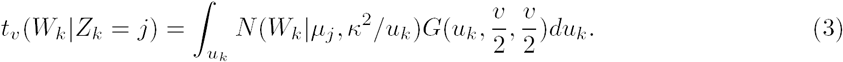

We also specify a normal distribution for logR, *W_k_*|*Z_k_* = *j* ∼ *N*(*μ_j_*, *σ*^2^) to examine how differently these two distributions of logR behave in terms of handling hypersegmentation.

We use the square of logOR due to the lack of the haplotype information ([16]). The squared logOR (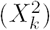) follows a chi-square distribution, 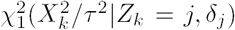 with one degree of freedom and a non-centrality parameter 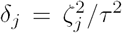, with the mean and variance *ζ_j_* and *τ*^2^, respectively. Finally, our *j*oint emission probability with a t-distribution of logR at the *k^th^* location given the state *j* is

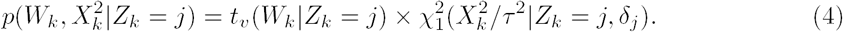

When logR follows a normal distribution, the first part on the right-hand side in Equation 4 is replaced by *N*(*W_k_*|*Z_k_* = *j*, *μ_j_*, *τ*^2^). Since logOR cannot be obtained in homozygous loci, the emission probability from Equation 4 is re-formulated, depending on the existence of logOR at the *k^th^* location (Equation 5).

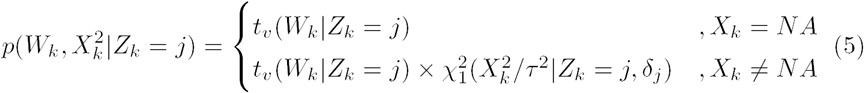

Here, *X_k_* = *NA* indicates that the *k^th^* location is homozygous, and heterozygous otherwise.

#### Estimation with E-M algorithm

The E-M (expectation-maximization) algorithm ([18]) is used to estimate the parameters of hsegHMM. In the E-step, the goal is to calculate the posterior probability, *P* (*Z_k_* = *j* | ***W***, ***X*^2^**) and the joint probability, *P* (*Z_k_* = *j*, *Z*_*k-*1_ = i| ***W***, ***X*^2^**), where ***W*** and ***X*^2^** indicate the sets of the logR and squared logOR values over chromosomes, respectively. These probabilities are evaluated by applying the forward-backward algorithm ([19]) resulting in the following conditional probabilities,

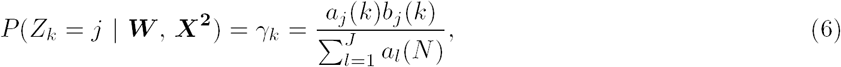

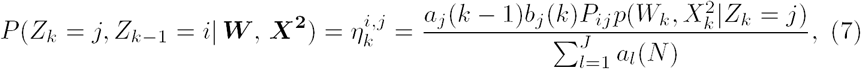

where 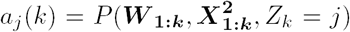 and 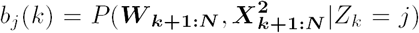; ***W*_l_:*m*** and 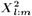 indicate the observed values of logR and squared logOR from the *l^th^* locus to *m^th^* locus. *a_j_*(*k*) is evaluated by a forward recursion up to the *k^th^* observation and the *b_j_*(*k*) by a backward recursion from the last to the (*k* + 1)*^th^* observation. The recursion equations are

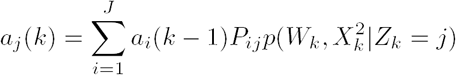

and

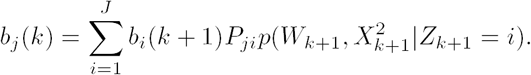

Then, the loglikelihood is simply computed as 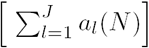. The calculation of *a_j_*(*k*) and *b_j_*(*k*) may result in extremely small values that cause an underflow issue, particularly with large *N* as in our application. Therefore, the scaled HMM [20] is implemented for all the analyses in this paper.

In the M-step, given the hidden state values obtained from Equations 6 and 7, we maximize the expectation of the conditional log-likelihood function with respect to all the parameters. The expectation of the complete log-likelihood function is given by,

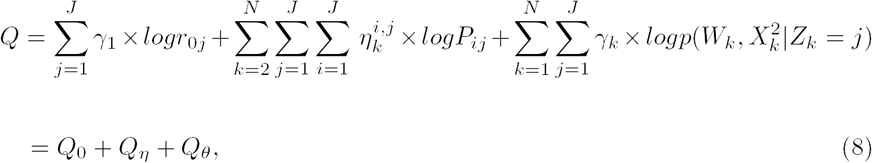

where *θ* is a set of global parameters: θ = { *α, Ψ*, *κ*^2^, *τ*^2^, *v* }. Given *γ_k_*and and 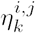 in Equation 8, we can estimate all the parameters by maximizing Q in the M-step. For estimating the initial probability *r*_0*j*_, we maximize *Q*_0_ under the constraint 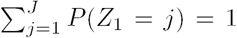,which is 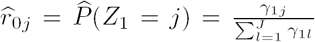. Similarly for estimating *P_ij_* under the constraint 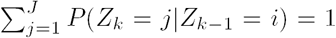, we obtain the closed form as

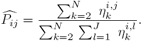

For estimating all the global parameters, we maximize Q*_θ_* in terms of them by using the L-BFGS-B optimization algorithm in the R function optim.

## Simulations

Initially, we perform two simulation studies for assessing how accurately our proposed model identifies true genotype states when hypersegmentation occurs.

For these two studies, we generate 500 datasets with true genotype sequences of size *N* = 10, 000 based on a Markov chain with a given transition probability, using the R- package markovchain. We consider four genotypes as true states: A, AA, AAAB, and AAAAB. The first two genotypes are chosen for hemizygous deletion and neutral LOH (loss of heterozygosity), while the last two are chosen for the amplification events. Moreover, AAAB (*j* = 7) and AAAAB (*j* = 10) give similar expectations for both logR and logOR, making it more challenging to distinguish between these two genotypes. In this study where *Ψ* = 1.6 and *α* = 0.9, *μ*_7_ and *μ*_10_ are 1.25 and 1.55 for logR; *ζ*_7_ and *ζ*_10_ are 1.03 and 1.31 for positive values of logOR. For each simulation study, the hsegHMM-N and the hsegHMM-T models are applied; The observed logOR*_j_* is generated by *X_k_* = *ζ_j_* +*ε_k_* where *ζ_k_* is normally distributed in these two simulation studies.

### t-distribution-based logR

In this simulation, we simulate hypersegmentation using a t-distribution for logR and examine both hsegHMM-N and hsegHMM-T models. We start with generating logR values from the t-distributions with *v*(= 4) degrees of freedom. The squared values of logOR are generated from the chi-square distribution with one degree of freedom. Similar to the TCGA-KL-8331 dataset used in the TCGA study section, we assign 90% of loci to be homozygous.

For each locus, the allele-specific SCNA is identified by choosing the genotype with the largest posterior probability. In order to evaluate the accuracy of our models, we estimate the probability of a correct identification across the genome. First, we create the classification index variable which is set to be 1 if the estimated genotype is correctly assigned for each locus in one simulation, and zero otherwise. We then calculate the probability of identification of each genotype across locus by averaging the index across 500 datasets. Figure 4 shows the probability of identification for all genotype states. It is obvious that the red lines (by the hsegHMM-T model) are significantly closer to the true signal (black line) than the blue lines (by the hsegHMM-N model) for all genotypes. In particular, the accuracy plot for the genotype AAAB shows much lower blue lines, compared with the red ones. For instance, the 5909*^th^* locus is truly assigned to genotype AAAB, and the probability of identification for AAAB at the location is 0.488 with the hsegHMM-N model but 0.940 with the hsegHMM-T model. This indicates that AAAB is harder to correctly identify when we use hsegHMM-N model rather than the correct hsegHMM-T model.

**Figure 1.**
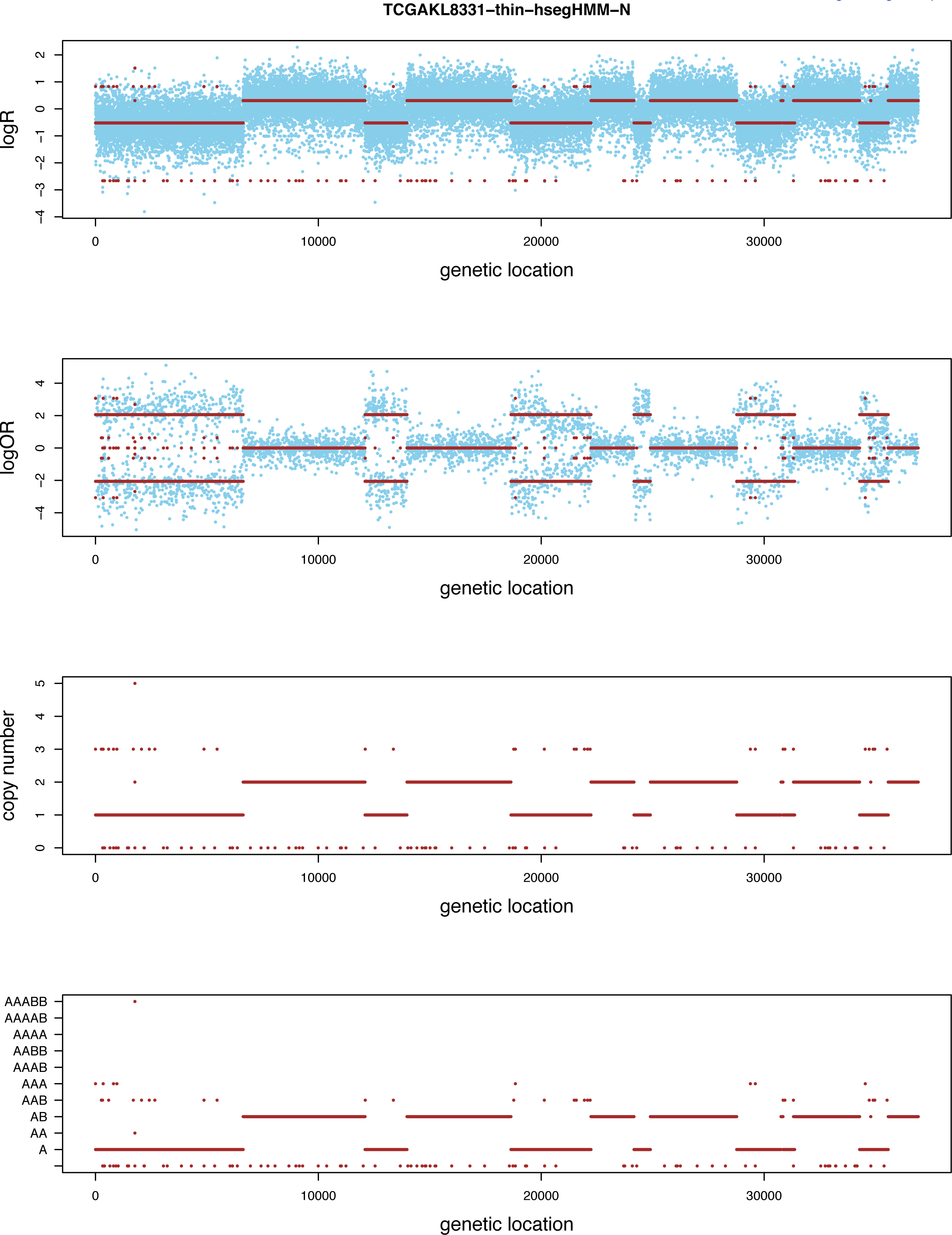
Allele-specific SCNA analysis based on the hsegHMM-N model of a renal cell carcinoma from a TCGA project (TCGA-KL-1883) The blue dots are observed values and red bars are estimates; The first two panels show the profiles of logR and logOR over the entire chromosomes; The last two panels indicate estimated copy numbers and genotype for each sequence over the entire chromosomes.

**Figure 2.**
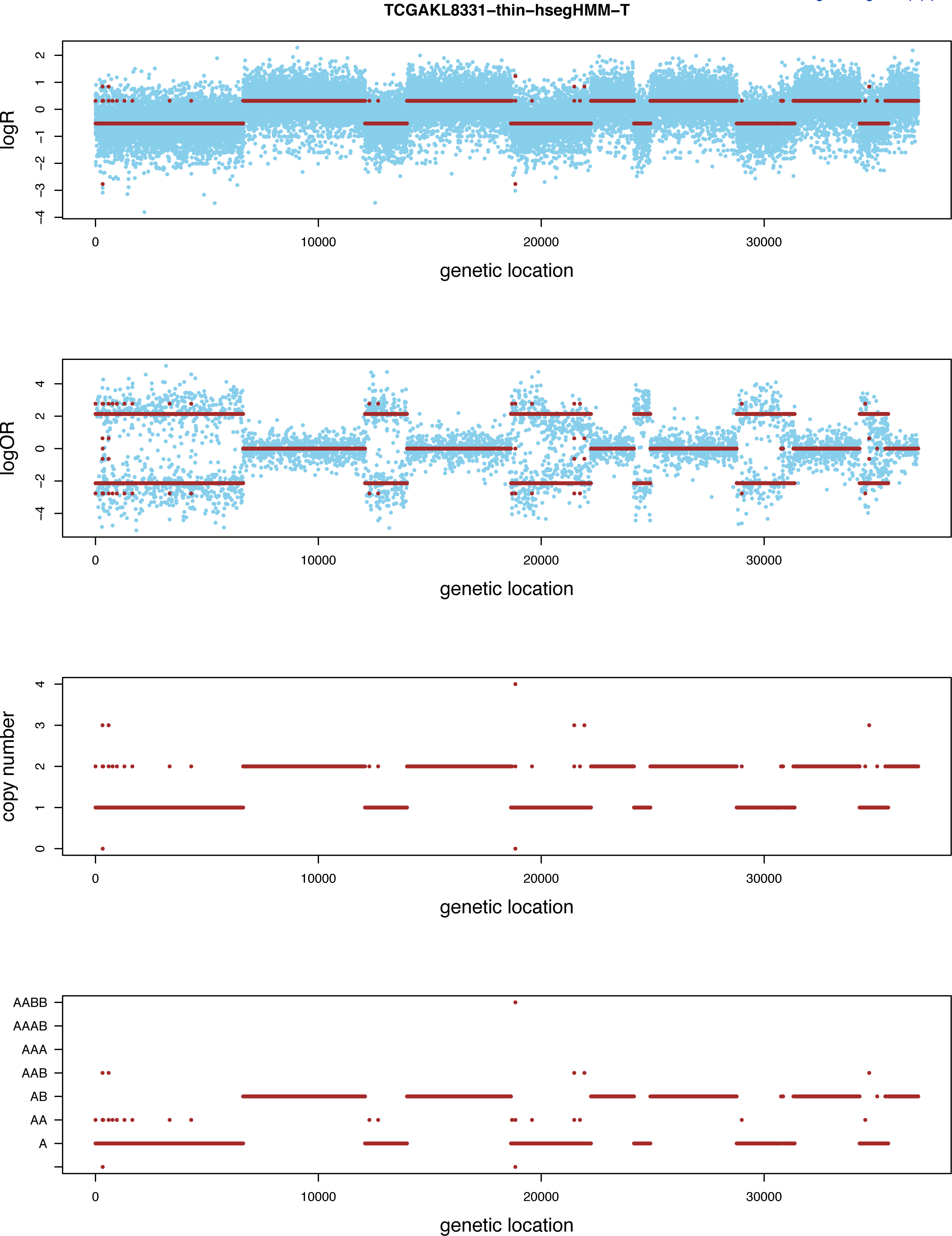
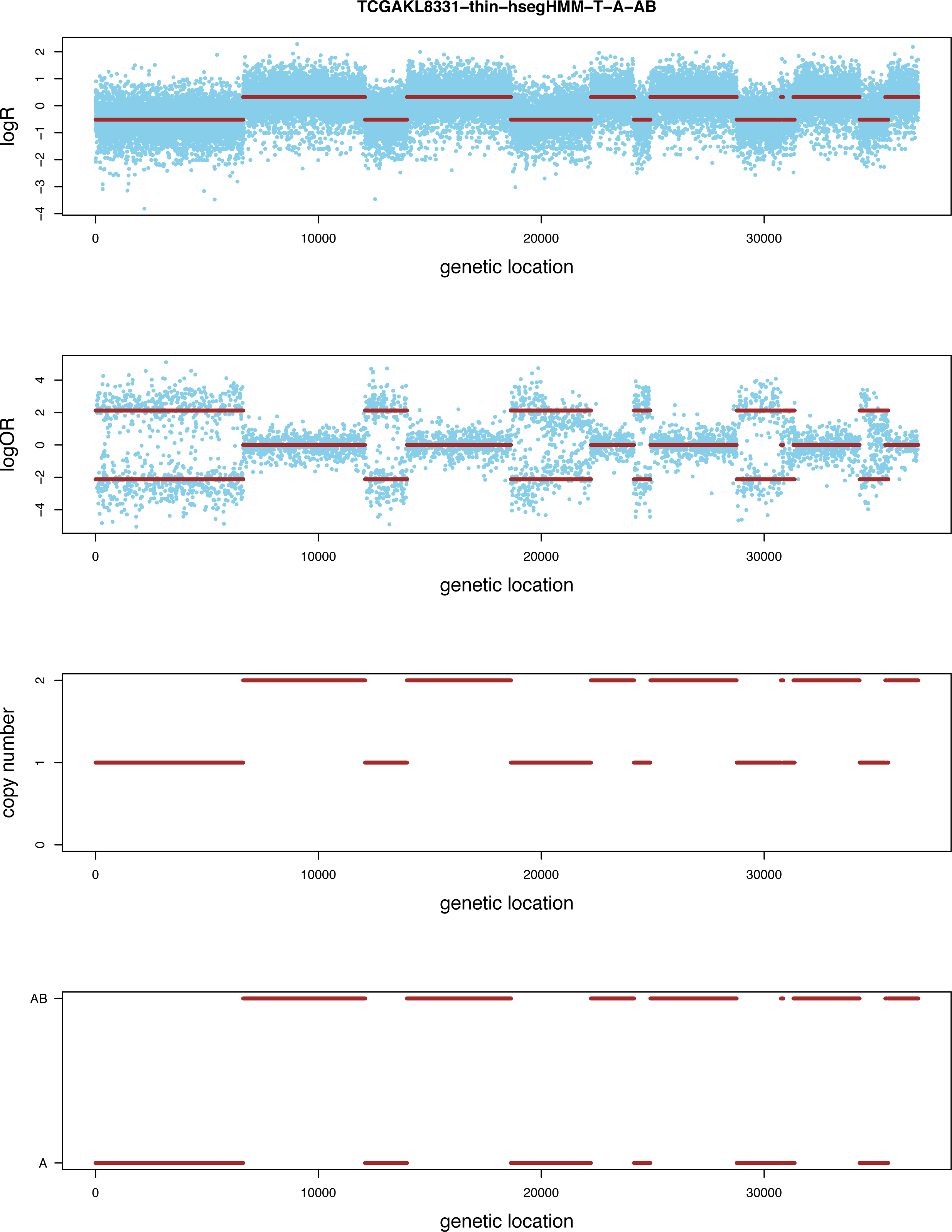
Allele-specific SCNA analysis based on the hsegHMM-T model (a) and the same model with the A and AB state space (b) of a renal cell carcinoma sample from a TCGA project (TCGA-KL-1883) The blue dots are observed values and red bars are estimates; The first two panels show the profiles of logR and logOR over the entire chromosomes; The last two panels indicate estimated copy numbers and genotype for each sequence over the entire chromosomes.

**Figure 3.**
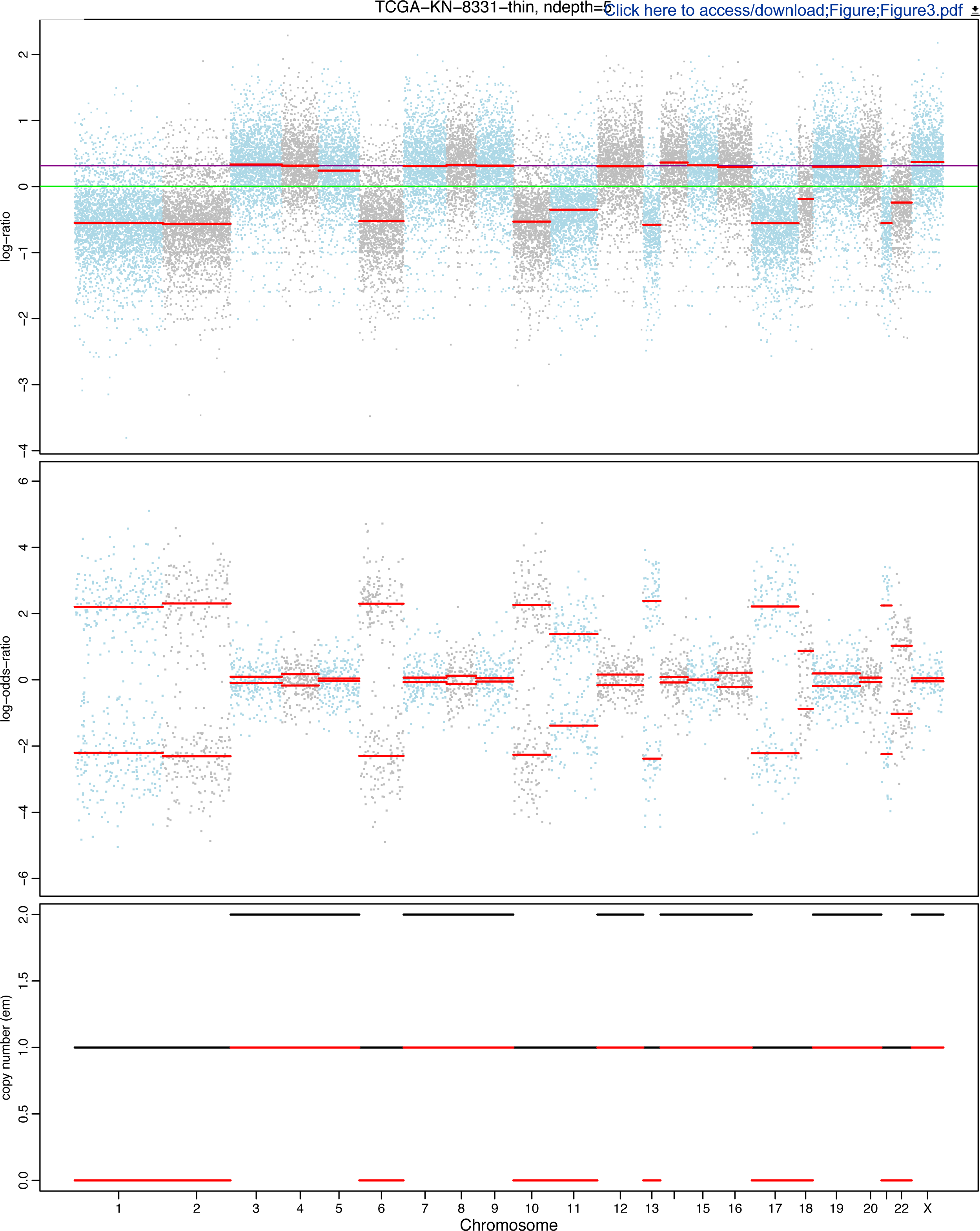
Allele-specific SCNA analysis based on the FACETS model of a renal cell carcinoma sample from a TCGA project (TCGA-KL-1883) The first two panels show the profiles of logR and logOR over the entire chromosomes; The last panel indicates estimated copy numbers of total and minor alleles (black and red lines, respectively) for each sequence over the entire chromosomes.

**Figure 4.**
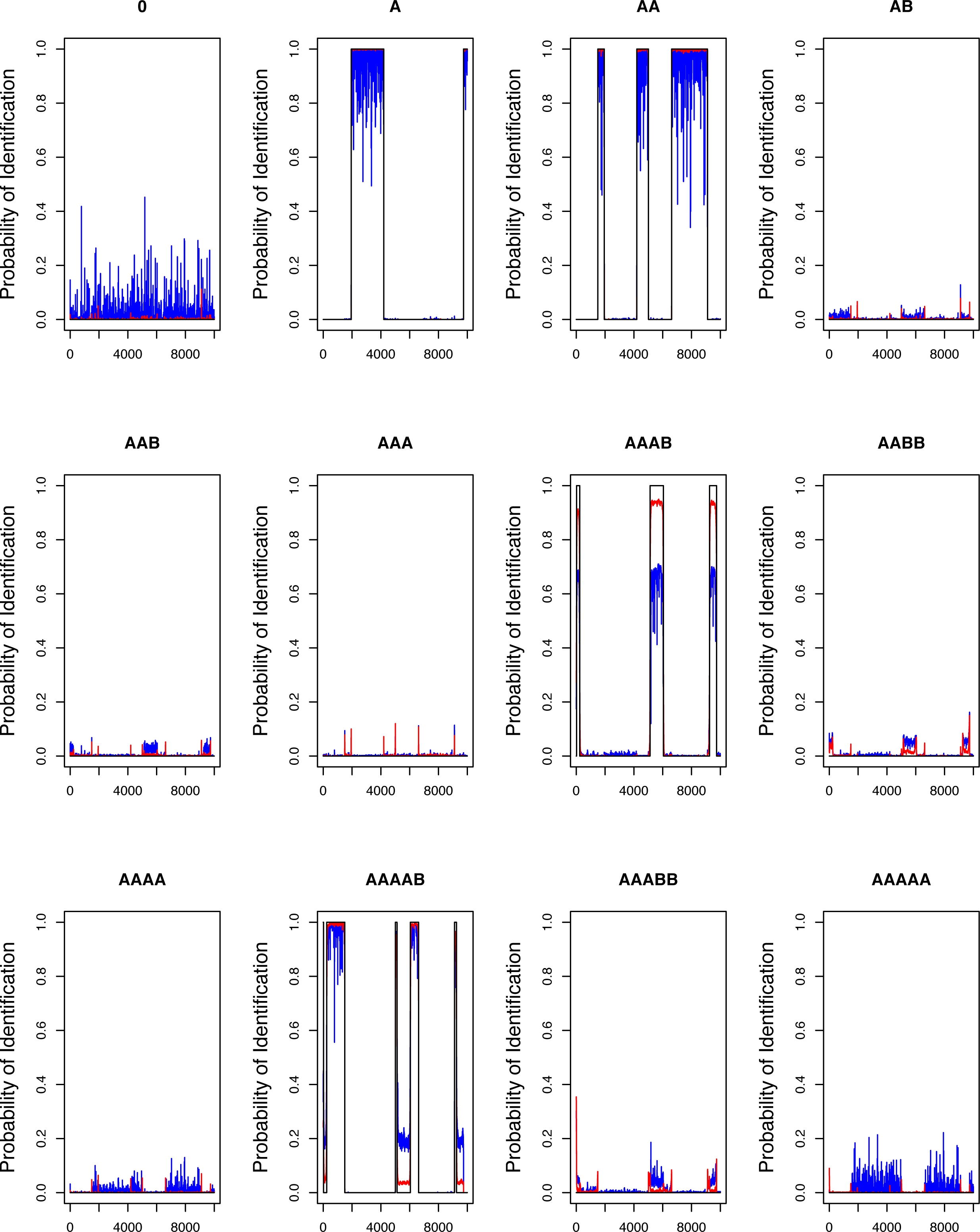
Probability of identification for all the genotype states from 500 simulated datasets with logR generated from the t-distribution. blue lines and red lines indicate the probabilities of identification based on the hsegHMM-N and hsegHMM-T models, respectively; Each dataset consists of 10,000 observations of logR and logOR.

We examine the statistical properties of the global parameter estimates in Table 3. SEs are empirical standard errors and SE_H_ are computed by averaging 500 asymptotic standard errors based on the Hessian matrices. The Hessian matrix for each dataset is numerically obtained by hessian in R package numDeriv. The estimates are unbiased even under the misspecified normal model. However, the asymptotic standard error estimates are sensitive to misspecification of the logR distribution (SE_H_ is different from SEs under the misspecified hsegHMM-N model).

**Table 3.**
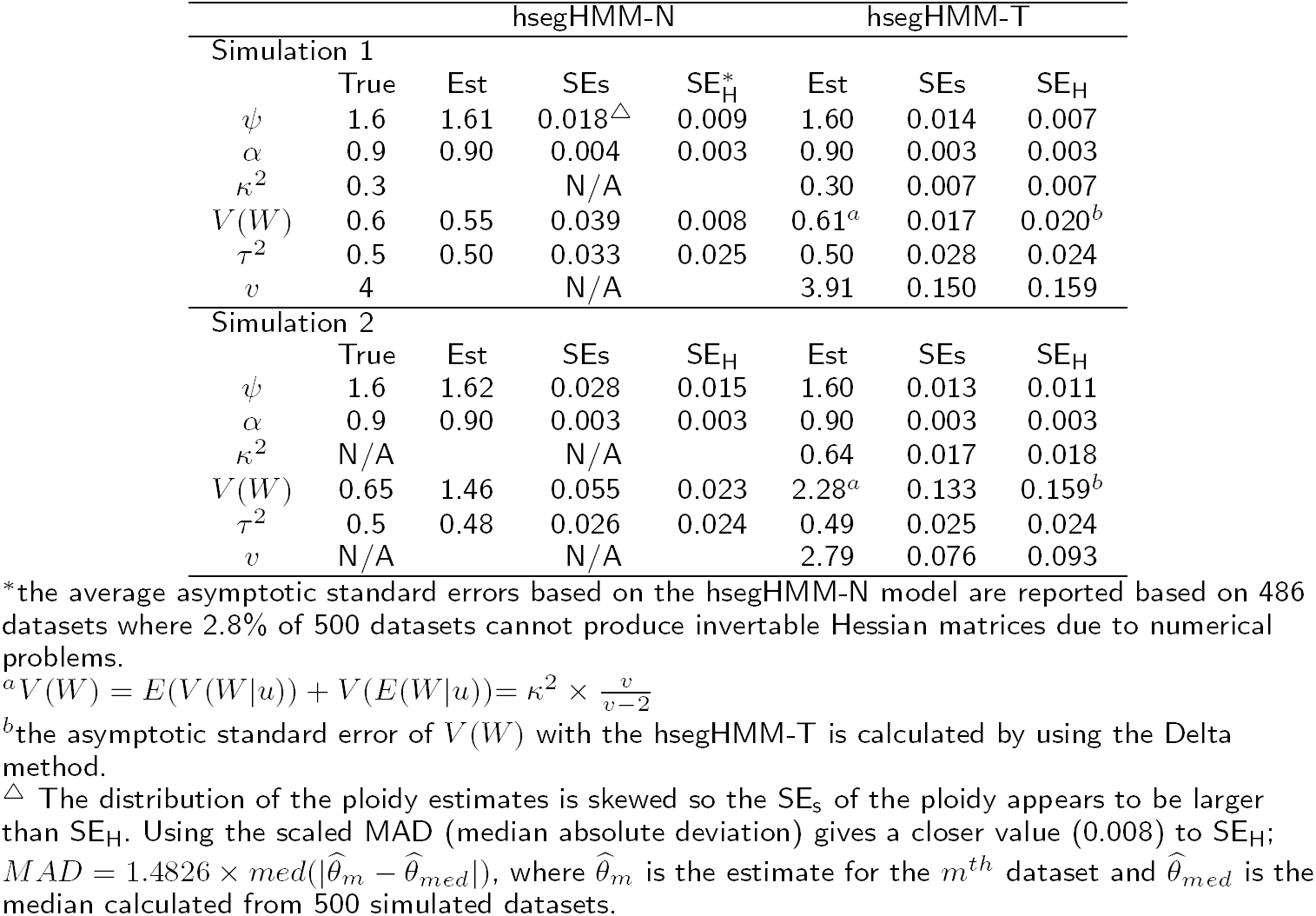
Summary of simulation studies with hsegHMM-N and hsegHMM-T models based on 500 simulated datasets. Simulation 1 and Simulation 2 are the t-distribution-based and the normal-mixture-based studies. Each dataset consists of 10,000 observations of logR and logOR. Est is average estimates from 500 datasets; *Ψ* is the ploidy, *α* is the tumor purity; *κ*^2^ is the variance component of logR in hsegHMM-T; V (W) and *τ*^2^ are the variance of logR and logOR in both models, respectively; SEs indicates the Monte-Carlo standard errors calculated from 500 datasets; SE_H_ indicates the average asymptotic standard errors of estimates based on the Hessian matrices.

### A mixture of normals-base logR

We examine the robustness of the t-distribution to alternative distributions that exhibit long tails. Specifically, we simulate under a mixture of normal distributions and examine the robustness of the hsegHMM-T model. Errors are generated by 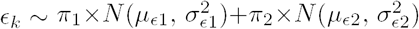, where *π*_1_ and *π*_2_ indicate the mixture proportions of the first and the second distributions, respectively. The means and variances for those two normal distributions are chosen under the condition of *E*(*∊_k_*) = 0 and 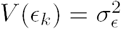. In this simulation, *π*_1_ and *π*_2_are set as 0.7 and 0.3 with *μ*_∊1_ = *μ*_∊2_ = 0,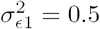, and 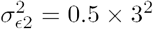. Then, the total error variance 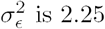.

Both the probability of identification (Figure 5) and the summary of estimators with corresponding standard errors, SEs and SE_H_ (Simulation 2 in Table 3) are shown in the same way as described in the previous Section (t-distribution-based logR). Figure 5 shows that all the red lines based on the hsegHMM-T model appear noticeably closer to the black lines than the blue lines based on the hsegHMM-N model for all the genotype states. In particular, detecting both AAAB and AAAAB with the hsegHMM-T model performs much better than the hsegHMM-N model.

**Figure 5.**
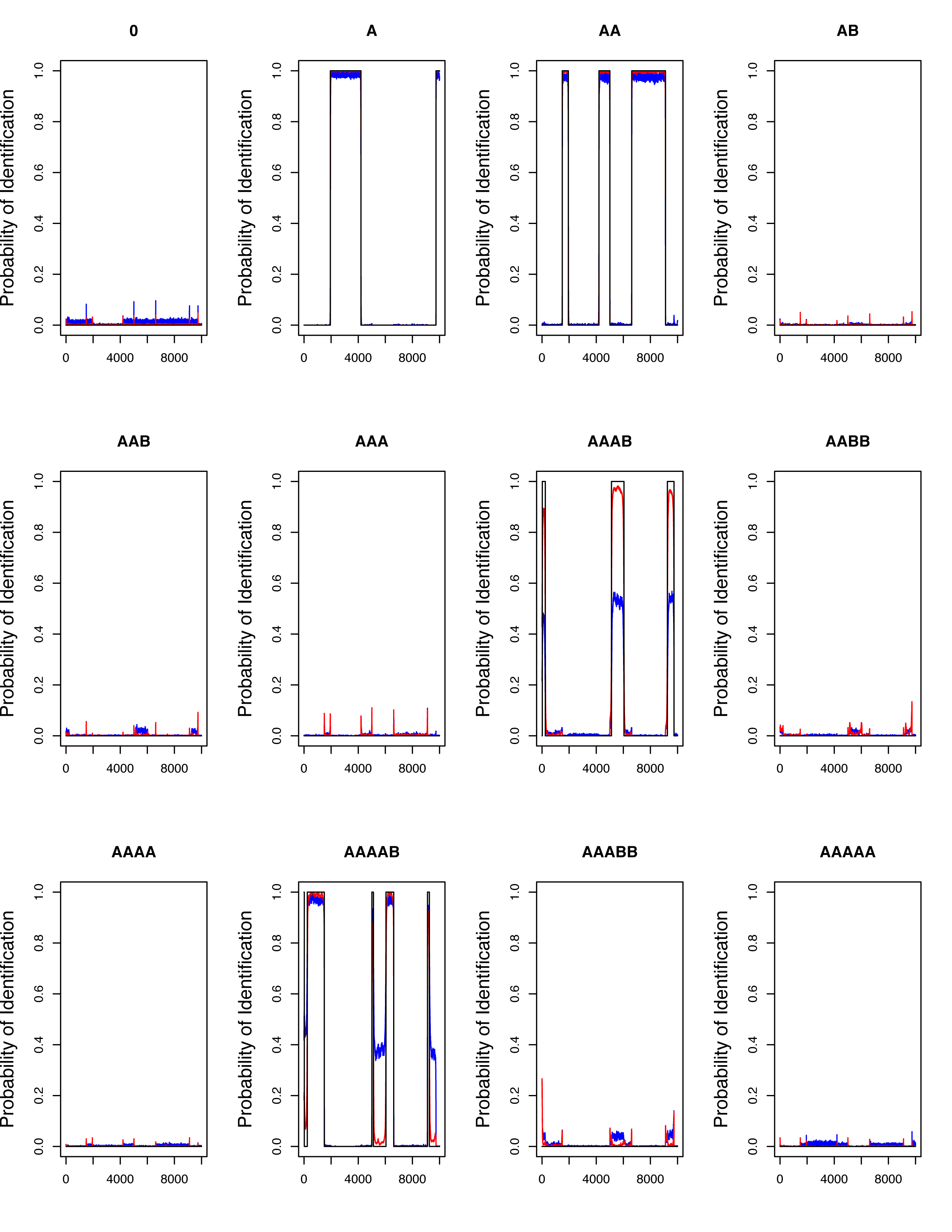
Probability of identification for all the genotype states from 500 simulated datasets with logR generated from the normal-mixture distribution. blue lines and red lines indicate the probabilities of identification based on the hsegHMM-N and hsegHMM-T models, respectively; Each dataset consists of 10,000 observations of logR and logOR.

The results indicate that under mild amounts of misspecification (t-distribution rather than a mixture of normals) of the log R distribution, estimates of global parameters and their standard errors are both unbiased, and the accuracy of classifications are very good. Consequently, hsegHMM-T provides more accurate estimates of genotype status by managing hypersegmentation more effectively than hsegHMM-N.

A single dataset-based result is provided to investigate more closely how much more robust the hsegHMM-T model is than the hsegHMM-N model to cope with hypersegmentation (Supplementary Figs S1-S4). The first two figures (Supplementary Figs S1 and S2) and the last two figures (Supplementary Figs S3 and S4) show the copy number profiles of a particular dataset from the first and second simulation scenarios, respectively. Similar to the analysis results, hsegHMM-T appears to handle hypersegmentation much better than hsegHMM-N. This same pattern was observed for all simulated datasets (data not shown).

We also perform additional simulation studies for different values of purity (*α* = 0.3, 0.5, 0.7) and different numbers of reads (half and double related to the original from Figure 6). Based on 500 simulated datasets, our proposed model performs better as the purity increases in terms of a higher probabilities of correct genotype identification (Figure S5).

**Figure 6.**
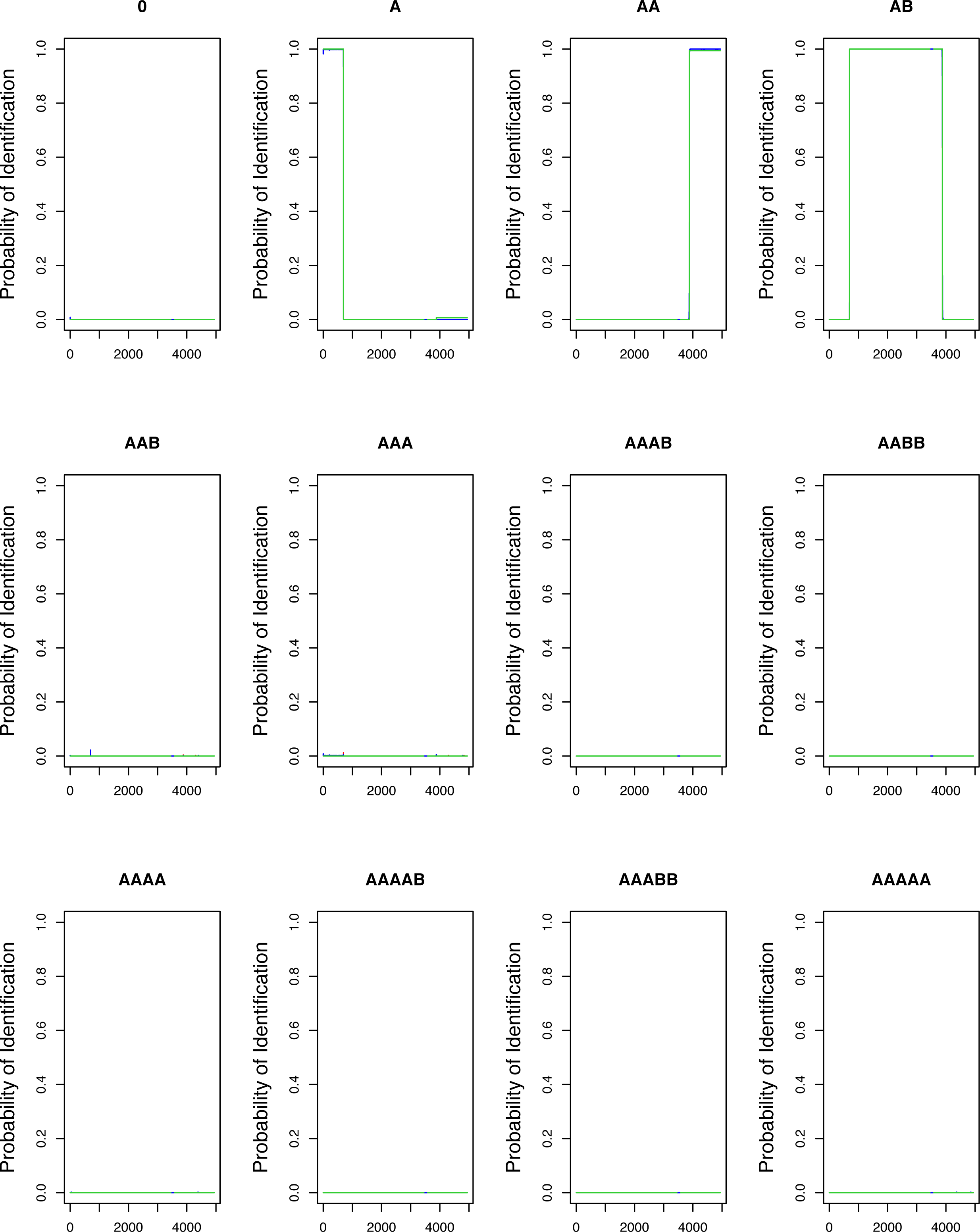
Probability of identification for all the genotype states from 500 simulated datasets based on creating read counts for normal and tumor cells. green lines, blue lines, and red lines indicate the probabilities of identification based on the FACETS, hsegHMM-N, and hsegHMM-T models, respectively; Each dataset consists of 4,942 observations of logR and logOR.

### A read counts-based simulation for the comparison of hsegHMM and FACETS

In this simulation, we compare our method, hsegHMM with FACETS which also constructs its model based on logR and logOR but with a segmentaion-based approach for allele-specific SCNA analysis. In order to make a fair comparison between these two methods, we generate datasets from read counts and read depth (coverage) in the beginning without relying on model assumptions of hsegHMM or FACETS. In this study, we choose the true profile of genotypes as A, AB, and AA orderly with different lengths over the entire chromosomes.

First, we consider the fact that the total coverages for normal cells and tumor cells are different. Thus, we assign different read depths for tumor and normal cells as 160 and 40 in this study. Then, for normal cells, the read depth is 40 and the read count of A allele is either 20 for AB (heterozygosity) or 40 for AA (homozygosity) regardless of the true genotypes for tumor. On the other hand, for tumor cells, both read depth and read counts are assigned depending on different genotypes. For the genotype AB, the read count and depth are set to be 80 and 160, respectively. For the genotype AA, the read count and depth are the same as 160. For the genotype A, the read count and depth are both 80 as the same number with a half of the total coverage since B allele is lost.

In order to generate read counts and depths, we use a uniform distribution with different intervals for normal and tumor samples. The intervals provide the variation occuring from measurement errors. For normal, read counts and depths are generated from a uniform distribution with the range of ±15 intervals for both the genotype AA and AB, and the range of ±20 for the genotype A. For tumor, read counts and depths are generated from either of these two uniform distributions with the range of ±30 and ±15 with probability of 70% and 30%, respectively. This setting provides logR and logOR values with different variances and asymmetric ranges between genotype A and the others, which makes more challenging to analyze. Finally, we round the decimal values of read counts and depths from these continuous uniform distributions to the nearest intergers.

Through the preprocedure of FACETS, we obtain total 4,942 values of logR and logOR based on the integer values of read counts and depths, for which both hsegHMM and FACETS are applied. Figure 6 shows how different these two methods behave through the probability of identification plots for all the different genotype states. These plots are based on 500 simulated datasets. hsegHMM-N (red lines) and hsegHMM-T (blue lines) have almost the exact patterns of the identification plots with FACETS (green lines) for all the genotypes. On the other hand, when the read depth distribution was skewed (a rescaled beta distribution with shape parameter values of 1 and 6), FACETS did poorly in region identification as compared to hsegHMM. Specifically, we consider both the cases of short and wide region length. Figure S6 shows the probabilities of correct identification for a short region based on 500 simulated datasets. hsegHMM-T identified the mutation in approximately 96% of the datasets as compared with 5% using FACETS (the left panel in Figure S6). For the wider region, FACETS improved relative to hsegHMM, but there was still a marked improvements of our approach (the right panel in Figure S6): 83% and 99% identification for FACETS and hsegHMM, respectively. These results show the advantage of hsegHMM as compared with FACETS for uneven coverage.

We also examine our method with different numbers of reads for read counts and depths by reducing half size and increasing double size of them (Figures S7 and S8). For both the half and double read size cases, our model shows similar results to those from the original size of reads (Figures S7 and S8). FACETS showed similar behavior when the read counts were altered (Figures S7 and S8).

## Results

### TCGA-KL-8331 renal cell carcinoma dataset

TCGA project (https://cancergenome.nih.gov) is a cancer genomic collaboration between the National Cancer Institute (NCI) and the National Human Genome Research Institute (NHGRI). This project includes critical genomic information of 33 types of cancers with more than two petabytes of TCGA genomic dataset to contribute cancer etiology, treatment, and diagnosis. In this research, we apply hsegHMM to whole-exome sequencing data from a chromophobe renal cell carcinoma (RCC) sample (TCGA-KL-8331).

TCGA-KL-8331 dataset consists of read counts and total depths for both normal and tumor paired tissues from the same patient over the entire chromosomes. This dataset contains 1, 217, 407 single nucleotide variants (SNVs). Through FACETS pre-processing step, these 1.2 MB SNVs were reduced to 369,131 SNVs, which are limited to the germline polymorphic sites and filtered by low quality including lower depth coverage positions (see the details in the Data pre-processing section in [16]). Thus, observed logR and logOR are calculated for *N* = 369, 131 loci. In this RCC sample, we find that approximately 13% (47,660) of loci are heterozygous with the corresponding logOR available. For computational feasibility, we perform a thinning process which keeps every 10*^th^* observation. This also reduces auto-correlations between observations, which helps alleviate hypersegmentation. We apply the hsegHMM procedure to the final dataset *N* = 36, 914.

Figures 1 and 2 show the results based on an assumed normal and t-distribution for the logR values given the hidden state (genotype state), respectively. We denote these two models as hsegHMM-N and hsegHMM-T, respectively. Each figure includes four panels corresponding to the estimated values of logR, logOR, copy numbers, and genotype status. With the hsegHMM-N model (Figure 1), estimated lines (brown color) show not only the main signals (longer bars) but also numerous dots across the chromosomes. These small dots occur due to the sensitivity of the hsegHMM-N model to extreme observations. The hsegHMM-T model reduces hypersegmentation with fewer short subsequences (Figure 2.(a)). However, a few numbers of short sequences still occur in using the hsegHMM-T model. Thus, instead of using the 12 genotype-state space, we consider only two major genotypes, A and AB identified by the hsegHMM-T with the 12 genotype states. It turns out that all the short dots are removed across the entire chromosomes (Figure 2.(b)). Thus, the hsegHMM-T model with the two major genotype states manages hypersegmentation most efficiently among those three different model fits. Furthermore, according to the model fitting criteria, the hsegHMM-T model with A and AB genotype states (hsegHMM-T_A/AB_) fits data best with the smallest AIC (62923.90) and BIC (62992.03). We also apply the FACETS method to compare the result with our method. The hsegHMM-T with the two major genotype states have almost the same allele-specific copy number profiles with FACETS in Figure 3.

The tumor sample purity *α* is estimated to be about 87-88% for all the methods, which indicates a high proportion of the tumor cells in the tumor tissue. The estimated ploidy, 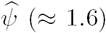 appears to be different from 2 in all the methods, which provides evidence for aneuploidy in this sequence. Note that the tumor purity, ploidy and variances of logR (V (W)) are similar in all the three hsegHMM models. This suggests that estimation of global parameters are robust to the distribution of logR and to an expanded genotype state space. This is in contrast to the allele specific genotype status that does appear to be sensitive to the distribution of logR and to the specification of an appropriate state space of genotypes, and hence to hypersegmentation.

## Discussion

We have shown that the hidden Markov modeling approach provides an effective way to identify allele-specific copy number alternations along the genome. As compared with FACETS, a segmentation-based approach, the hsegHMM provides an assessment of the uncertainty in parameter estimate (i.e. ploidy and purity), using likelihood-based estimate of variances as well as the ability to assess variability in copy number identification by computing posterior estimates of the genotype at each locus. It is also important to mention that hsegHMM is based on the output of WES, which in turn relies on the exome enrichment platforms where capture efficiency may still affect SCNA estimation.

A major focus of the paper was demonstrating that hypersegmentation in allele-specific SCNA data can be substantially reduced by incorporating a long-tailed emission distribution (hsegHMM-T model) into a HMM framework. We also found that hypersegmentation could occur by choosing a state space (possible genotypes) that is more expansive than necessary. Thus, we recommend that the most parsimonious model with a limited number of genotype states be chosen. Of course, the choice of this model should be based on using penalized likelihood methods such as AIC. Last, the hsegHMM assumes that logR and logOR measurements given genotype are independent across the entire chromosomes. This may not be true when loci are very close together, and failure of this assumption may lead to hypersegmentation. We therefore recommend thinning the sequence data (e.g., taking only one out of ten data points) to avoid this problem.

The application of hsegHMM can be extended in three future directions that have important applications in cancer genetics. First, hsegHMM can be applied to a population-based study where many subjects will be analyzed. In this case, we suggest that individual-specific analyses be conducted and the results combined in a final analysis. For example, evidence of a SCNA being related to a particular cancer may be suggested if a sizable proportion of the posterior probabilities of a genotype at a particular chromosome location are greater than a certain threshold (e.g. > 80%). Second, the relationship between a genetic factor and a subject-specific covariate may be examined in a second stage regression. For example, by using all the individual ploidy estimates obtained from the population-based study, we can construct a linear regression of the log ploidy estimate, 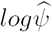 with a set of any covariates such as 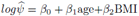. As an illustration for a population-based study, we have analyzed all 316 renal cell carcinoma samples from TCGA with the proposed model based on 5 copy number of state-space. We obtained the distribution of estimated ploidy across all the samples for any major copy number alteration event across the chromosomes. We also estimated the distribution of purity which is a measure of the quality of the tissue samples. Furthermore, we created a cytoband-based stacked histogram of allele-specific SCNA events for integrating allele-specific SCNA profiles from all the 316 samples. Each sample has its own allele-specific SCNA profile with different genotypes and regions. To standardize genetic locations across the samples, we used a cytoband file format which has predefined positions of cytobands across the whole chromosomes. For each cytoband, the corresponding allele-specific SCNA event is assigned within individual sample, and the number of times each event occurs is counted. After counting the frequencies of all the cytobands, we found that the most frequent mutation is a hemizygous deletion (genotype A) that has highest frequency on Chromosomes 3 and 14. In addition, we found a high frequency of a Gain (genotype AAB) in the region between q21.3 and q35.3 on Chromosome 5 (Figure S9).

Last, our model structure can be extended to infer tumor subclonal populations. In practice, a tumor sample contains a mixture of clones not just one main clone which is assumed in hsegHMM. Such an approach can also be embedded into a hidden Markov modeling framework, and is the subject of future research.

## Conclusions

In this paper, we propose a hidden Markov model framework (hsegHMM) for estimating genotype status as well as copy number at each locus, incorporating the complexities of tumor samples as well as hypersegmentation. Specifically, under certain type of data with more fluctuated or irregular observations, hsegHMM-T model performs better than hsegHMM-N model in terms of such a remarkable reduction of hypersegmentation. As a byproduct of the hsegHMM estimation procedure, we can compute the posterior probabilities of allele-specific genotype status (the Method section) as well as provide a rigorous comparison of different models (e.g. normal versus t-distribution) by using AIC and BIC (the Result section). Hence, hsegHMM provides a rigorous framework for statistical inference and model assessment. hsegHMM can also expand the genotype state space so that it can handle a more flexible range of copy number alterations. Specifically, this flexibility is useful for analyzing data from certain type of cancers with high-level amplification events. Simulation studies showed that hsegHMM-T performed much better than FACETS in situation where the coverage (read depth) is uneven across the genome.

In conclusion, hsegHMM offers an allele-specific SCNA analysis robust to hypersegmentation while accounting for tumor purity and ploidy. Such robustness enhances the accuracy of detecting genotype status at each locus in NGS-based platforms.

## Acknowledgements

We would like to acknowledge our usage of the data from The Cancer Genome Atlas (TCGA) supported by the National Cancer Institute and National Human Genome Research Institute: https://cancergenome.nih.gov. We would like to thank Bill Wheeler (Information Management Services) and Lei Song (Biostatistics Branch) for computational contributions.

## Funding

The work was supported by the Intramural Research Program of US National Institutes of Health, National Cancer Institute. This work utilized the computational resources of the NIH HPC Biowulf cluster. (http://hpc.nih.gov).

## Availability and Implementation

All the source codes of hsegHMM are available at https://dceg.cancer.gov/tools/analysis/hseghmm.

## Author’s contributions

HCW, PSA and B*Z* all contributed to the model formulation, analysis, simulations, interpretation, and writing. All authors read and approved the final manuscript.

## Ethics approval and consent to participate

Not applicable.

## Consent for publication

Not applicable.

## Competing interests

The authors declare that they have no competing interests.

## Author details

^1^Biostatistics Branch, Division of Cancer Epidemiology and Genetics, National Cancer Institute, National Institutes of Health, 20892 Bethesda, MD, U.S.A.. ^2^Biostatistics Branch, Division of Cancer Epidemiology and Genetics, National Cancer Institute, National Institutes of Health, 20892 Bethesda, MD, U.S.A..

## Additional Files

Figure S1 – Allele-specific SCNA analysis based on the hsegHMM-N model of a simulated dataset for the simulation study with logR generated from t-distribution

The first two panels show the profiles of logR and logOR over the entire chromosomes; The last two panels indicate estimated copy numbers and genotype for each sequence over the entire chromosomes.

Figure S2 – Allele-specific SCNA analysis based on the hsegHMM-T model of a simulated dataset for the simulation study with logR generated from t-distribution

The first two panels show the profiles of logR and logOR over the entire chromosomes; The last two panels indicate estimated copy numbers and genotype for each sequence over the entire chromosomes.

Figure S3 – Allele-specific SCNA analysis based on the hsegHMM-N model of a simulated dataset for the simulation study with logR generated from normal-mixture distribution

The first two panels show the profiles of logR and logOR over the entire chromosomes; The last two panels indicate estimated copy numbers and genotype for each sequence over the entire chromosomes.

Figure S4 – Allele-specific SCNA analysis based on the hsegHMM-T model of a simulated dataset for the simulation study with logR generated from normal-mixture distribution

The first two panels show the profiles of logR and logOR over the entire chromosomes; The last two panels indicate estimated copy numbers and genotype for each sequence over the entire chromosomes.

Figure S5 – Probability of identification for different sizes of purity with logR generated from the normal-mixture distribution

The red lines, green lines, and gold lines represent the high (*α* = 0.7), medium (*α* = 0.5), and low (*α* = 0.3) purity cases. All the results are conducted with hsegHMM-T.

Figure S6 – Probability of identification for a region generated from a non-standard beta-based read depths The red line and blue line represents hsegHMM-T and FACETS; The black dotted line is the true one.

Figure S7 – Probability of identification with the half size of read counts and depths from Figure 6 green lines, blue lines, and red lines indicate the probabilities of identification based on the FACETS, hsegHMM-N, and hsegHMM-T models, respectively; Each dataset consists of 4,942 observations of logR and logOR.

Figure S8 – Probability of identification with the double size of read counts and depths from Figure 6 green lines, blue lines, and red lines indicate the probabilities of identification based on the FACETS, hsegHMM-N, and hsegHMM-T models, respectively; Each dataset consists of 4,942 observations of logR and logOR.

Figure S9 – Frequency of Allele-specific SCNA events based on cytobands across all the chromosomes for 316 samples from TCGA

“HOMD” indicates homozygous deletion state.

